# LRSomatic: a highly scalable and robust pipeline for somatic variant calling in long-read sequencing data

**DOI:** 10.64898/2026.02.26.707772

**Authors:** Robert A. Forsyth, Luuk Harbers, Amber Verhasselt, Ana-Lucía Rocha Iraizós, Sidi Yang, Joris Vande Velde, Christopher Davies, Nischalan Pillay, Laurens Lambrechts, Jonas Demeulemeester

## Abstract

**Motivation:** Long-read sequencing is increasingly used in cancer research and clinical genomics due to its ability to resolve complex genomic variation and previously inaccessible regions of the genome. However, dedidated workflows for comprehensive somatic variant analysis from long-read whole-genome data remain scarce, limiting uptake in cancer genomics.

**Results:** We present LRSomatic, a Nextflow-based, nf-core-compliant pipeline supporting somatic SNV, indel, structural variant, and copy number calling from PacBio HiFi and ONT data. LRSomatic supports paired tumor-normal and tumor-only designs, as well as integration of epigenetic integration via Fiber-seq. Benchmarked on COLO829 and HG008 reference cell lines, LRSomatic achieves state-of-the-art performance across both platforms and variant types. Applied to a case of clear cell sarcoma, it recovers all identified driver alterations, including the pathognomonic EWSR1::ATF1 fusion, and resolves haplotype-specific chromatin accessibility via Fiber-seq.

**Availability and Implementation:** Freely available at https://github.com/intgenomicslab/lrsomatic, implemented in Nextflow DSL2, supported via Docker and Singularity.

## Background

Whole genome sequencing (WGS) has revolutionized genetic analysis in both research and clinical practice. Comprehensive detection of genomic variation at base pair resolution has allowed researchers to capture novel genomic drivers and risk variants associated with diseases such as Alzheimer’s disease, cancer, coronary artery disease, and a myriad of rare disorders^1–5^. In clinical practice, WGS has been used successfully to identify therapeutically actionable variants in cancer, screen for heritable genetic disorders, and detect rare diseases for early treatment in newborn care^6–9^. Due to the wealth of information contained within WGS data, dedicated bioinformatics workflows have become essential to efficiently and reproducibly detect (relevant) variation. This has led to the usage of both dedicated bioinformatics workflow management systems, like *Nextflow*, and standardized good practice guidelines, like *nf-core*, to ensure portability and reproducibility in WGS analysis^10,11^.

Within this ecosystem, open-source pipelines have been developed for reliable identification of germline and somatic variation from WGS data, such as *Oncoanalyser* and *Sarek*^12,13^. However, these pipelines currently support only short-read sequencing data. While short-read WGS remains a powerful tool for variant discovery, particularly in the case of single nucleotide variants (SNVs) and small insertions and deletions (Indels), the approach struggles to resolve complex alterations, such as structural variants (SVs), and those located in highly polymorphic (e.g. *HLA*), recently duplicated, or repetitive regions of the genome (8–15% of the genome)^14–17^. Long-read sequencing methods, such as Oxford Nanopore Technologies (ONT) nanopore and Pacific Biosciences (PacBio) HiFi sequencing overcome these limitations by readily generating reads with a median length of ≥10 kilobases, allowing for comprehensive resolution of all types of variation^9,18^. Although pipelines exist for identifying germline variation using long reads, such as *Nallo*, there are few analysis workflows for detecting somatic variation in a platform-agnostic manner ^19^.

When applied to native DNA samples, both PacBio and ONT sequencing can also detect endogenous base modifications such as CpG (5mC) methylation, providing insights into genome function. This ability to simultaneously provide genetic and epigenetic insights can be further extended to include chromatin accessibility patterns through techniques like Fiber-seq^20^. Fiber-seq uses a nonspecific DNA N^6^-adenine methyltransferase to stencil positions in chromatin accessible regions, allowing researchers to resolve regulatory architectures genome wide^20^. While this technique can be used to infer functional implications of genomic variation, currently only few specialized analytical tools exist which can effectively integrate and exploit the combined genetic and epigenetic information^21^.

To address the shortage of workflows for long-read whole-genome somatic analysis and boost functional insights via inclusion of endogenous and exogenous base modification information, we developed *LRSomatic*, a *Nextflow*-based bioinformatics workflow^22^. Built on *nf-core* best practices for reproducibility and portability, *LRSomatic* is technology agnostic, reliably identifying somatic variation from PacBio or ONT sequencing data, while incorporating optional epigenetic information from base modifications. By unifying these capabilities into a single analytical framework, *LRSomatic* enables researchers to extract a comprehensive picture of genomic and epigenomic variation at base pair resolution.

## Implementation

### Overview

*LRSomatic* is implemented in *Nextflow DSL2*, a specialized workflow management system optimized for bioinformatic analyses^10^. This platform provides native support for multiple containerization systems, like Docker and Singularity, along with diverse execution environments, including SLURM, Amazon AWS, PBS, and Google Cloud Platforms^10,23–25^. This architecture ensures both reproducibility and portability across computational infrastructures.

Moreover, *LRSomatic* adheres to *nf-core* guidelines, guaranteeing standardized modular tool usage and comprehensive documentation. Designed with flexibility as a core principle, *LRSomatic* supports both PacBio HiFi and ONT sequencing platforms and accommodates various experimental designs including paired tumor-normal and tumor-only samples. *LRSomatic* also incorporates modules for epigenetic data capture obtained through technology-specific 5mC and 6mA calling. A schematic overview of the workflow for each data type is given in **Fig 1**.

**Figure 1:**
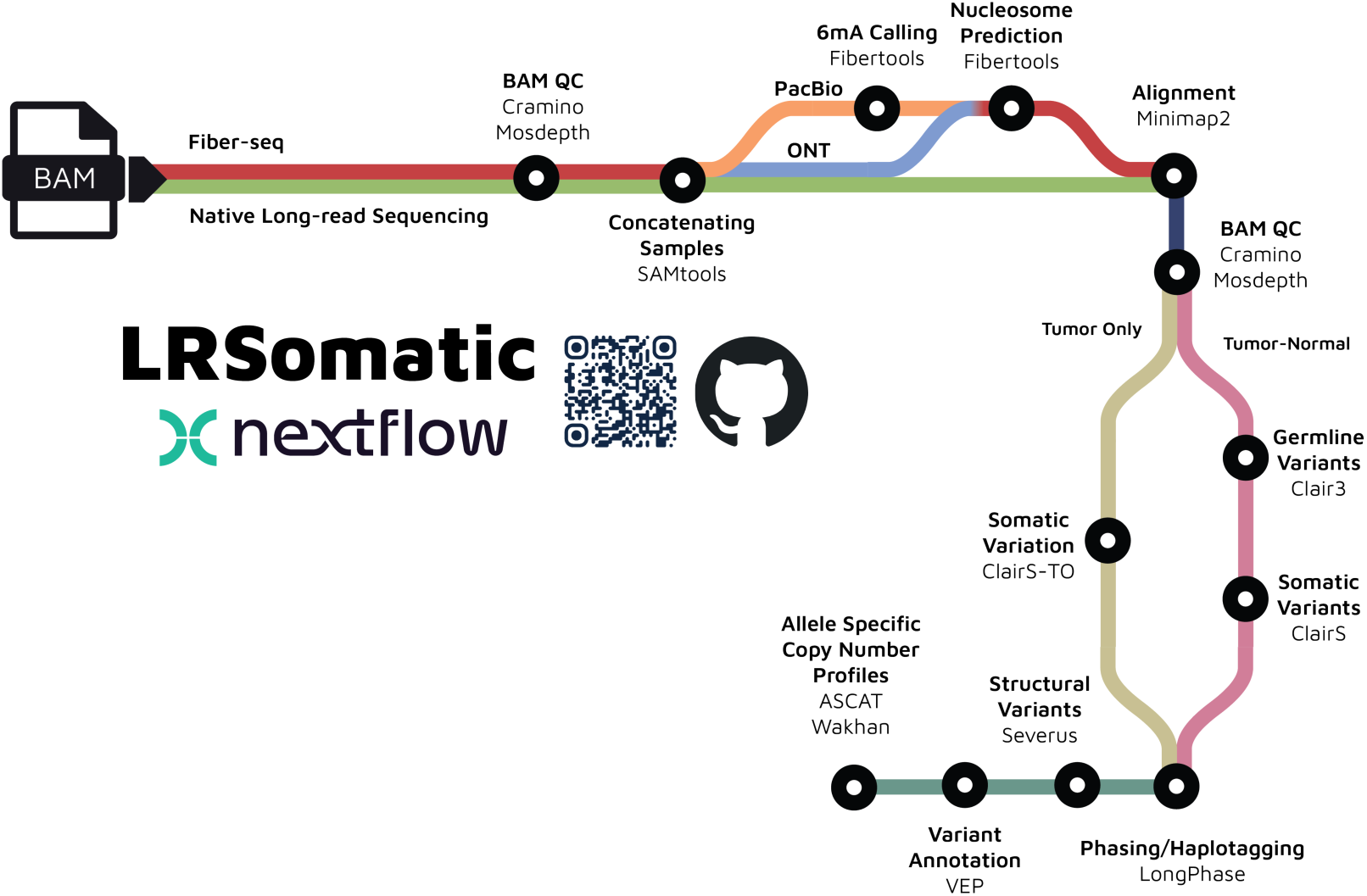
Metro map style overview of LRSomatic, including key module functionalities and tools.

### Initial Processing and Alignment

*LRSomatic* expects a comma-separated data spreadsheet with sample names, paths to basecalled unaligned BAM files from samples, along with sample-specific information about sex, Fiber-seq usage, and sequencing platform. The pipeline will concatenate BAM files associated with a single sample to ensure biological replicates are processed together. Quality control (QC) metrics are extracted from each BAM file using *Cramino*, a QC tool designed specifically for long-read sequencing data^26^.

Subsequently, Fiber-seq samples are processed through the *Fibertools* suite of modified basecalling and nucleosome prediction tools^27^. Accurate 6mA prediction is essential for interpretation of accessible chromatin regions in Fiber-seq^20^. Optional prediction of 6mA modification is performed for reads from PacBio HiFi read. This step is included to maintain compatibility with older PacBio output containing kinetics data, as 6mA calling can be done on-device using recent versions of the software (Instrument Control Software (ICS) v13.3 on Sequel II and Revio; Jasmine ≥2.4.0). 5mC calling is not included in *LRSomatic*. ONT sequencing data, as opposed to PacBio HiFi data, requires modified base inference to be coupled with initial basecalling. Since basecalling falls outside the scope of this pipeline, only optional *Fibertools* 6mA prediction for PacBio reads is included. Following 6mA prediction, both PacBio and ONT reads undergo downstream nucleosome and regulatory element prediction.

Next, reads are aligned to a chosen reference genome through *Minimap2*^28^. *LRSomatic* natively supports GRCh38 and T2T-CHM13 (hs1) reference genomes, however any custom reference genome can be used^14^. GRCh38 represents the latest assembly of the Genome Reference Consortium and boasts currently unmatched annotation resources and tool support. Use of the assembly version without ALT contigs is recommended for long-reads. However, there are large gaps across current GRCh38 assemblies, particularly in repetitive, centromeric, and subtelomeric regions^14^. In order to ensure users can perform variant calling in these regions, *LRSomatic* has support for all tools to perform analysis using the more recent telomere-to-telomere complete T2T-CHM13v2.0 reference genome. Once alignment is complete, a final quality control analysis is performed on the aligned BAMs via *Cramino*^26^.

### Variant Calling and Phasing

Post-alignment, samples are separated based on the presence of a matched normal control. Paired tumor-normal samples have germline variants called by *Clair3*, a tool which harnesses both a pileup and full alignment model to call SNVs and Indels^29^. Somatic variants are then identified using *ClairS*, a variant caller built on a similar methodology but tuned to discriminate somatic and germline variation by taking into account the intricacies of tumor-derived data, such as copy number alterations, subclonal variation, and normal cell admixture^30^. For tumor-only samples, *ClairS-TO* is employed for both germline and somatic small variant calling^31^. Specifically designed for tumor-only sequencing data, *ClairS*-TO leverages two neural networks, one to predict the likelihood a variant is somatic and one to predict the likelihood that the variant is not^31^.

Germline SNVs and Indels are subsequently phased and used as a basis for haplotagging reads using *Longphase*, a state-of-the art algorithm for phasing and haplotagging long-read sequencing data^32^. The resulting haplotagged BAM files and phased germline variants are then leveraged to identify structural variants using *Severus*, which employs haplotype-aware breakpoint graphs to model complex rearrangements^33^. Next, *ASCAT* and *Wakhan* utilize SNP loci data to perform allele-specific copy number calling, tumor purity and ploidy inference, with *Wakhan* additionally incorporating phased breakpoint information^34,35^.

All variants receive comprehensive functional annotation through Ensembl’s Variant Effect Predictor (VEP)^36^. Each sample is, by default, run with ‘--everything’ command, ensuring that each variant is labeled with each curated database available to VEP^36^. This includes ClinVar’s prediction of variant pathogenicity and connections to disease and treatment, COSMIC’s annotation of somatic variation associated with cancer, predicted effects on the resulting proteins from amino acid changes via SIFT and PolyPhen2, among other extensive VEP resources^36–40^.

## Methods

### Sample Selection

To demonstrate *LRSomatic’s* capabilities, we analyzed existing data from COLO829 and HG008 cell lines, as well as in-house data from a case of Clear Cell Sarcoma (CCS15). We downloaded sequencing data from COLO829 melanoma and matched lymphoblastoid cell lines from both PacBio and ONT public repositories^41,42^. We additionally obtained both PacBio and ONT sequencing data for the HG008 pancreatic ductal adenocarcinoma cell line and its matched pancreatic normal tissue from the Genome in a Bottle Consortium (GIAB)^43^. COLO829 and HG008 PacBio samples were sequenced on a PacBio Revio and those from ONT were sequenced on PromethION R10.4.1 flow cells (v14 chemistry) with a 5kHz sampling rate^41–43^. CCS15 metastatic and matched normal muscle tissue were processed for Fiber-seq on both PacBio and ONT and also subjected to lllumina short-read sequencing, as detailed below.

### Short-read sequencing

Frozen material from the Royal National Orthopaedic Hospital biobank (London, UK) was cryo-sectioned and examined by a pathologist to ensure >50% tumor content and <20% necrosis or inflammation. DNA was extracted from tumor and adjacent muscle curls using the QIAamp DNA Mini kit (Qiagen, 51304) and quality controlled using nanodrop, Qubit and Tapestation to ensure high purity and integrity of samples for sequencing. Tumor and matched normal DNA were prepared using the TruSeq PCR-free kit and paired-end sequenced (2×150bp) on an Illumina NovaSeq 6000 to a minimum whole-genome coverage of 70X and 30X, respectively.

### Nuclei isolation & open chromatin labeling

Fresh-frozen CCS15 biopsy samples were sectioned into 10–15 curls of 20 µm thickness using a cryostat. All curls were collected in a 1.5 mL Eppendorf tube, and 500 µL of ice-cold nuclei extraction buffer (10 mM Tris-HCl, pH 7.4; 10 mM NaCl; 0.1% BSA; 0.5 mM spermidine, pH 7.4; 0.1% Tween-20; 0.02% digitonin) was added. The mixture was transferred using a 1 mL wide-bore tip to a pre-chilled KIMBLE dounce tissue grinder (Sigma-Aldrich, D9063) and homogenized on ice with 10–20 strokes of pestle A. The homogenate was incubated for 5 min on ice and subsequently quenched by addition of 1 mL ice-cold wash buffer (10 mM Tris-HCl, pH 7.4; 10 mM NaCl; 0.1% BSA; 0.5 mM spermidine, pH 7.4; 0.1% Tween-20). The suspension was filtered through a 100 µm cell strainer into a 2 mL round bottom Eppendorf tube, and nuclei were pelleted by centrifugation at 300×g for 5 min at 4°C. Supernatant was removed, and the nuclear pellet was washed twice by resuspension in 500 µL wash buffer followed by re-centrifugation at 300×g for 5 min at 4°C. Finally, nuclei were filtered through a 40 µm cell strainer, collected in a 2 mL Eppendorf tube, counted and quality checked using a LUNA-FL Automated Fluorescence Cell Counter.

For 6mA labeling, 1 × 10⁶ nuclei per sample were pelleted by centrifugation at 300×g for 5 min at 4°C. The pellet was gently resuspended using wide-bore tips in 200 µL Fiber-seq labeling buffer (15 mM Tris, pH 8.0; 15 mM NaCl; 60 mM KCl; 1 mM EDTA, pH 8.0; 0.5 mM EGTA, pH 8.0; 0.1% BSA; 0.5 mM spermidine, pH 7.4), freshly supplemented with 0.8 mM S-adenosylmethionine (NEB, B9003S) and 3.5 µM recombinant Hia5 N^6^-adenine methyltransferase^20,44^. The mixture was incubated in a thermomixer for 10 min at 25°C at 800 rpm. The reaction was stopped by proceeding directly to DNA extraction using the Monarch High Molecular Weight (HMW) DNA Extraction Kit for Tissue (NEB, T3060).

### Fiber-seq PacBio Library prep

HiFi sequencing libraries were prepared following the manufacturer’s instructions for the PacBio SMRTbell prep kit 3.0 (102-166-600) with the SPRQ chemistry. Briefly, 2 µg of extracted, 6mA-labeled HMW DNA from tumor and matched normal CCS15 samples was diluted in 130 µL low TE buffer and sheared to ∼20 kb using a Megaruptor 3 (Diagenode, B06010003; shearing syringes E07010003) at speed setting 31. Sheared DNA was purified using a 1X SMRTbell cleanup and eluted in 47 µL of elution buffer. End repair and A-tailing, adapter ligation and cleanup, and nuclease treatment were performed according to the SMRTbell prep kit 3.0 protocol. Size selection was performed using a PippinHT system (Sage Science) with a 0.75% gel cassette (HPE7510). Elution was done for 30min starting at 12 kb to yield DNA fragments between 12 and 40 kb in size. Size-selected DNA underwent a 1X SMRTbell cleanup, eluted in 25 µL elution buffer prior to performing annealing, binding and cleanup as per the Revio SPRQ polymerase kit protocol (103-520-100). Final libraries were sequenced on a single Revio SMRT Cell each (PN 102-202-200).

### Fiber-seq ONT Library prep

ONT libraries for Fiber-seq were prepared using either the Native Barcoding Kit 24 V14 (SQK-NBD114.24; ONT; CCS15 matched normal sample) or the Ligation Sequencing Kit V14 (SQK-LSK114; ONT; CCS15 tumor sample). Briefly, 1.5 µg of 6mA-labeled HMW DNA was diluted in 130 µL low TE buffer and sheared on a Megaruptor 3 (Diagenode B06010003; shearing syringes E0700003) with speed setting 25. Sheared DNA was purified using 1.5X AMPure XP beads (A6388; Beckman Coulter) and eluted in either 11 µL (NBD114.24) or 47 µL (LSK114) Elution Buffer. The eluted DNA served as input to library preparation as per the ONT kit protocols. Final libraries were each sequenced for 72 h on a PromethION 24 using 1 flow cell loaded with 50 fmol.

### Pipeline Processing

We processed all PacBio and ONT samples through *LRsomatic* in both tumor-normal and tumor-only modes using GRCh38 without ALT contigs as the reference. The Illumina sequencing data from CCS15 was processed using *Oncoanalyser*, a gold standard *nf-core* implementation of the WiGiTS pipeline by the Hartwig Medical Foundation designed to comprehensively analyze short read sequencing data from cancer samples^45^. As a critical aim of both pipelines is to identify (clinically relevant) somatic driver mutations, we used *Oncoanalyser’s* calls as a benchmark for *LRSomatic’s* performance on CCS15.

## Results

### Time and Memory Usage

Complete processing of the six samples across both tumor-normal and tumor-only workflows required 3,570 CPU hours on a high-performance compute cluster (Flemish Supercomputing Center). Nodes were equipped with 2x Intel Xeon Platinum 8360Y CPUs@2.4 GHz (IceLake; 36 cores each) and 256GB RAM. For the full sample set, *LRSomatic* generated a 1.9 TB output directory alongside a 5.4 TB working directory, with sample-specific disk usage detailed in **Table 1**. Intermediate BAM files were the primary driver of storage consumption in both output and working directories across all samples. ONT BAM files exhibited greater post-alignment size increases compared to PacBio files, resulting in proportionally higher disk usage relative to their input size. Moreover, CCS15 PacBio data contained polymerase kinetics information, which was stripped after being used for ***6mA*** prediction in the pipeline, leading to smaller subsequent intermediate BAM files.

**Table 1:** Summary of Input BAM file sizes and disk usage of files directly associated with the sample in working and output directories. Note: this excludes reference files required across multiple runs, e.g., reference genome (indexed) fasta file.

| Sample Name | Tumor input<br>BAM size | Normal input<br>BAM size | Work Directory<br>Disk Usage | Output<br>Directory Disk<br>Usage |
| --- | --- | --- | --- | --- |
| CCS15-ONT<br>(paired) | 68 GB | 95 GB | 838 GB | 248 GB |
| CCS15-ONT<br>(tumor-only) | 68 GB | N/A | 450 GB | 99 GB |
| CCS15-PacBio<br>(paired) | 320 GB | 305 GB | 609 GB | 131 GB |
| CCS15-PacBio<br>(tumor-only) | 320 GB | N/A | 375 GB | 64 GB |
| COLO829-PacBio<br>(paired) | 63 GB | 58 GB | 270 GB | 132 GB |
| COLO829-PacBio<br>(tumor-only) | 63 GB | N/A | 195 GB | 61 GB |
| COLO829-ONT<br>(paired) | 167 GB | 137 GB | 814 GB | 409 GB |
| COLO829-ONT<br>(tumor-only) | 167 GB | N/A | 516 GB | 165 GB |
| HG008-PacBio<br>(paired) | 61 GB | 51 GB | 238 GB | 116 GB |
| HG008-PacBio<br>(tumor-only) | 61 GB | N/A | 190 GB | 61 GB |
| HG008-ONT<br>(paired) | 118 GB | 44 GB | 508 GB | 235 GB |
| HG008-ONT<br>(tumor-only) | 118 GB | N/A | 478 GB | 153 GB |

Plots of per-task duration, CPU usage (as percentage of a single core), maximum memory usage, and CPU utilization are shown in **Fig. 2**. The most substantial contributors to computational demand include *Fibertools* prediction of 6mA, germline variant annotation, and the Clair suite (*Clair3*, *ClairS*, and *ClairS-TO*). These resource-intensive steps present several opportunities for optimization. Utilizing on-device 6mA calling for PacBio HiFi (Jasmine; available since ICS v13.3 update) can significantly reduce both pipeline duration and memory usage. The Clair tools additionally offer an optional “fast” mode that decreases runtime and memory requirements, albeit with reduced prediction confidence^29^. Moreover, both Clair suite tools and *Fibertools* are equipped with GPU support which can reduce runtime significantly. Finally, germline annotation, or any other non-essential module, can be skipped to further minimize computational demands based on specific analysis needs.

**Figure 2:**
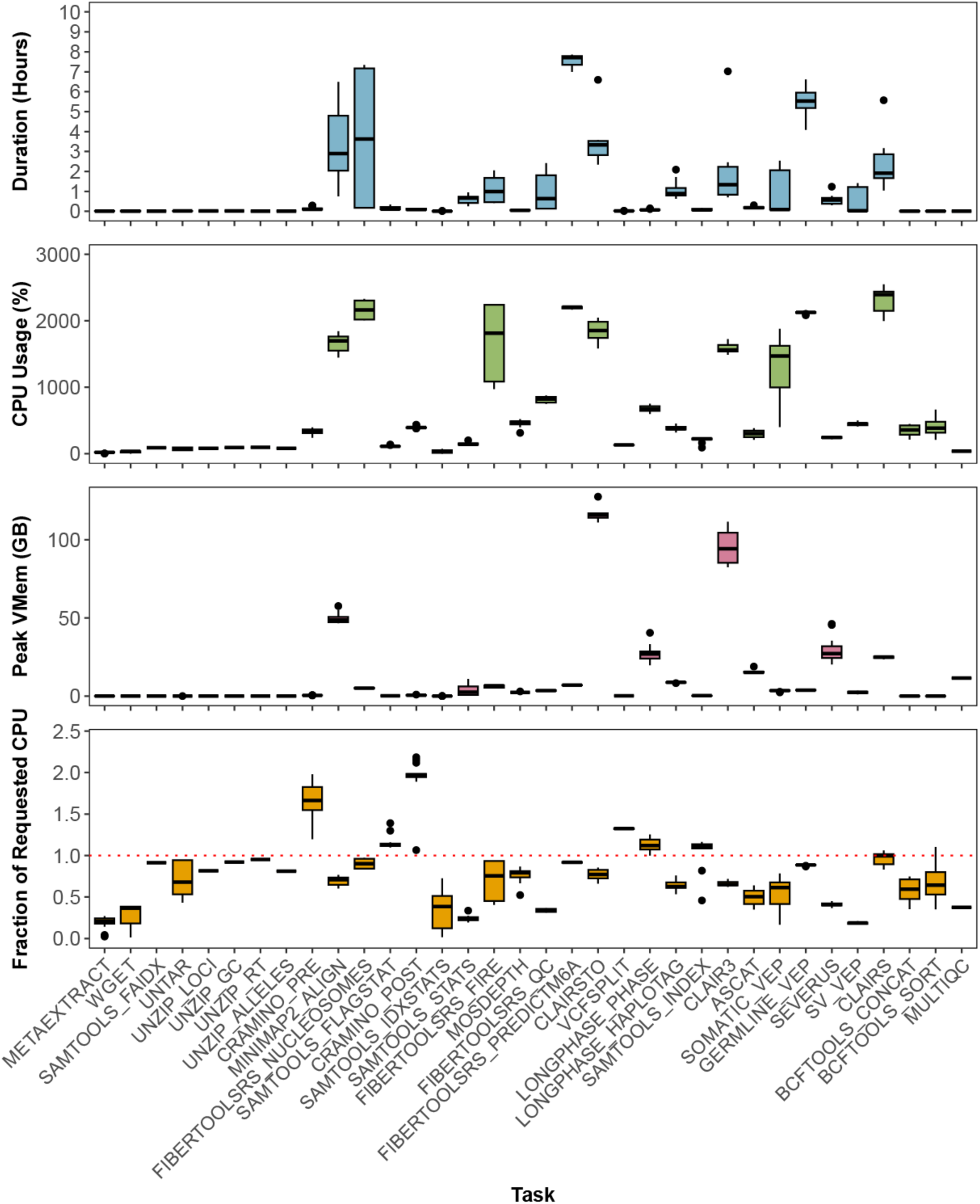
Summary of per task run statistics for each sample run. From top to bottom: Duration per task, CPU usage per task given as a percentage of a single core, Peak Virtual Memory usage per task (RAM + Disk swap), CPU utilization given as a fraction of requested task CPUs (red line given at 100% utilization for clarity).

Despite these resource intensive steps, all tasks remain within reasonable computational bounds. Even the most demanding tasks were completed using less than a single node, making this pipeline accessible to most computational frameworks. Requested resources are used efficiently with each task using a median of 82% of its requested CPUs. Storage utilization is similarly efficient: on average, each sample generated 3.53× (range: 2.13 - 6.62) its input size in intermediate files and 1.22× (0.97 - 1.52) its input size in output files. These summary statistics exclude PacBio data for CCS15, where kinetic data stripping reduces intermediate storage requirements leading to misleadingly small ratios of input to intermediate file sizes. Furthermore, inclusion of kinetic data is less representative of current PacBio sequencing output which now supports on device modification calling. Collectively, *LRSomatic* demonstrates efficient use of both compute and storage resources, highlighting its practical utility as a comprehensive somatic analysis pipeline.

### Quality Control Metrics

After alignment to GRCh38, samples had a median aligned data yield of 113.06 Gbp [61.80; 138.63], a median coverage of 35.06X [19.90; 36.41], and a median N50 read length of 17,365 base pairs [12,748; 35,084) (**Fig. 3**). The GIAB ONT sequenced HG008 normal sample had notably lower total yield and coverage. As the corresponding tumor sample has high coverage, somatic variation can still be robustly detected by tools in *LRSomatic*, but some bleed through of germline variants may be expected. Reads from CCS15 Fiber-seq sequencing on PacBio had a median length of 14,066 bp, with reads containing a median of 1078 6mA and 70 5mC calls. From the ONT Fiber-seq samples, reads possessed a median length of 8053 base pairs, with each read containing a median of 262 6mA and 54 5mC calls. *Fibertools* predicted a median of 66 nucleosomes and 35 nucleosomes per read from PacBio and ONT technologies respectively (**Fig. 4**).

**Figure 3:**
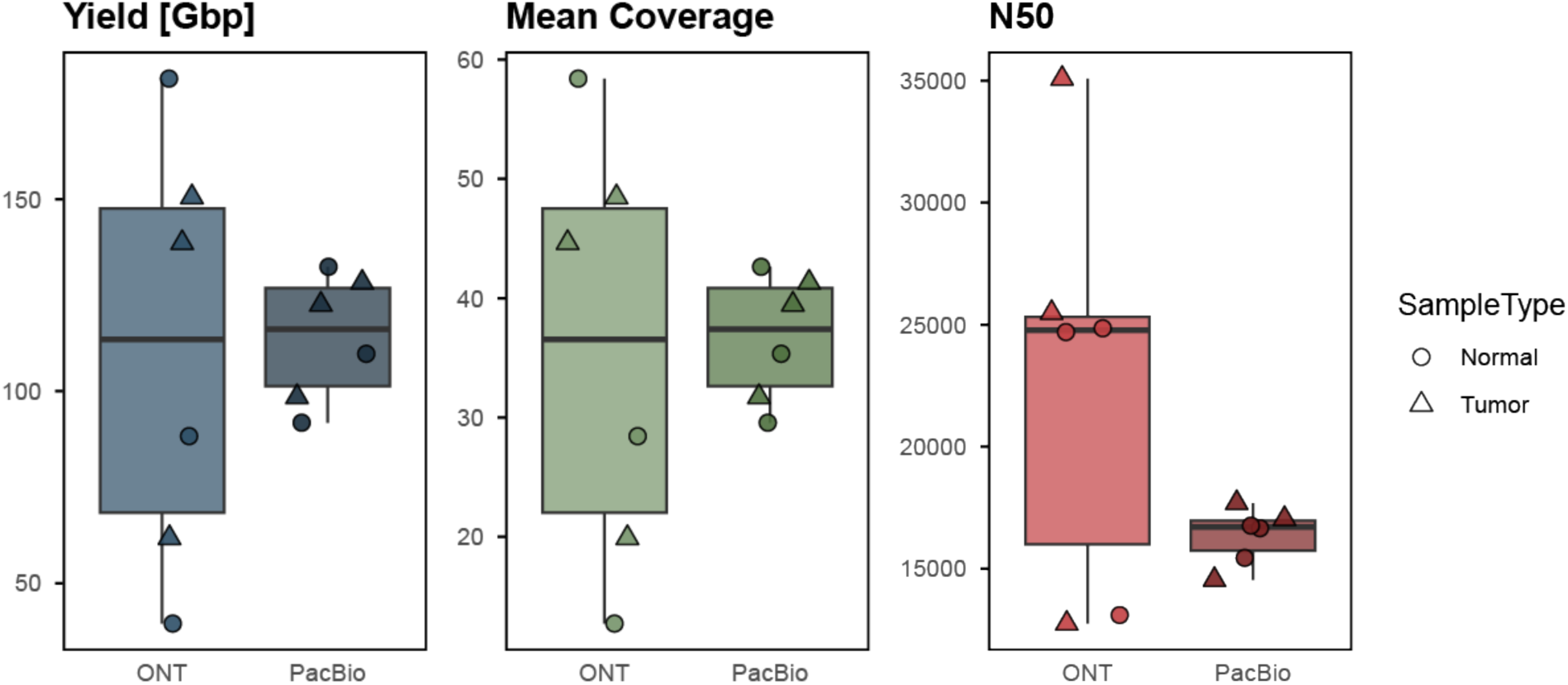
Boxplots (from left to right) of yield (in Gigabases, Gbp), mean coverage, and read length N50 for all samples separated by sequencing technology.

**Figure 4:**
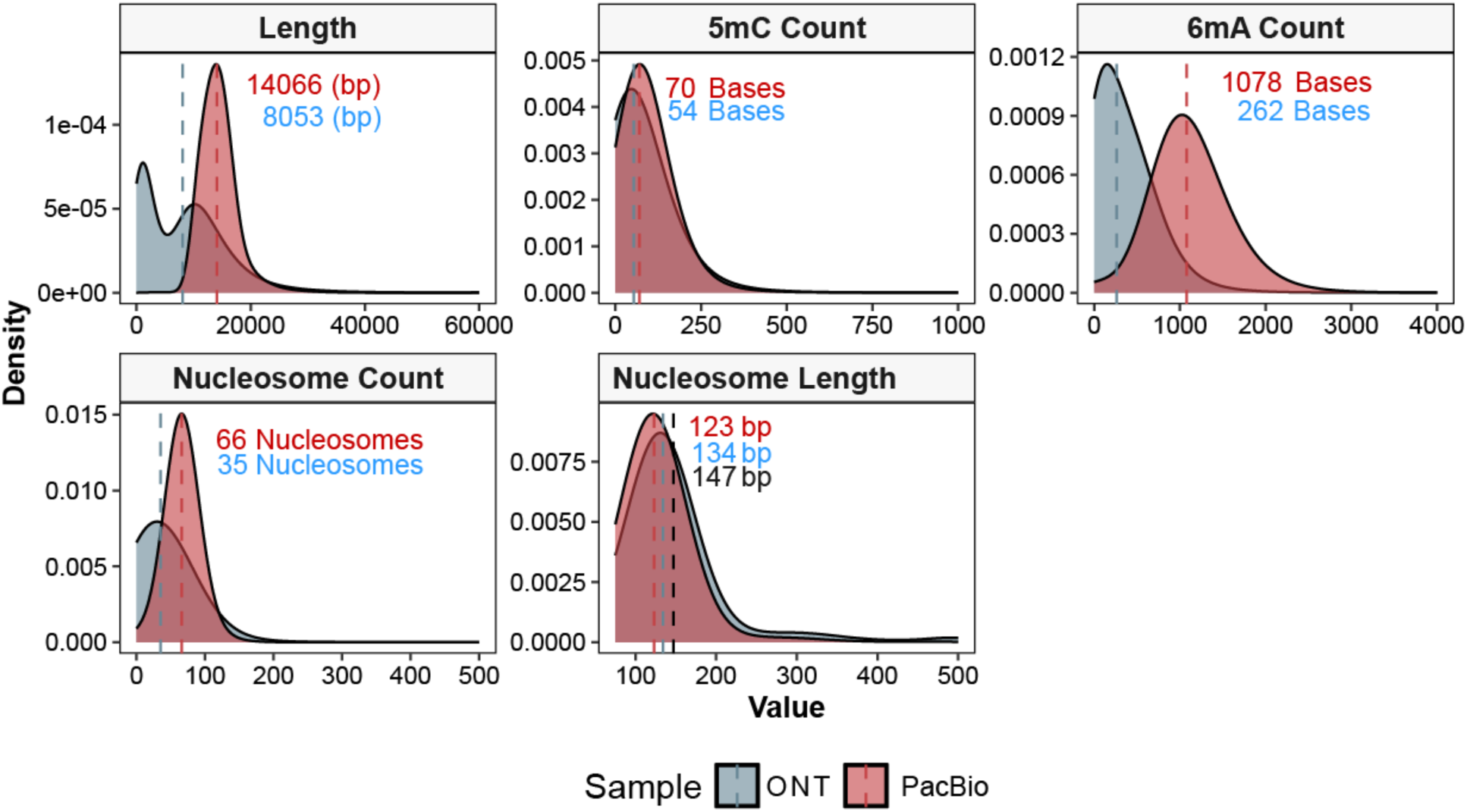
Kernel density plots of CCS15 Read Length (top left), 5mC Count (top middle), 6mA Count (top left), Nucleosome Count (bottom left), and predicted Nucleosome Length (bottom middle) separated by sequencing technology. Median values for PacBio and ONT distributions are marked with dashed lines with exact values labeled on the graph. The length of a human nucleosome (147 bp) is marked in the nucleosome length plot.

### Somatic Small Variant Calling

To confirm *LRSomatic’s* ability to recover small somatic variants, we compared the *ClairS* and *ClairS-TO* calls to an established reference set on COLO829 inferred by *MuTect2*, *Strelka2*, *Lancet* (exonic regions), and *Manta* (indels only) on high coverage (up to 278x) short-read sequencing data (Illumina HiSeqX and NovaSeq)^46^. Variant calls were included in this ground truth set if they were marked as High Confidence, identified by at least 2 callers, had a variant allele frequency (VAF) >0.05, and their position was covered by at least 4 reads in both the tumor and normal. This filtered somatic set contained 36,421 SNVs and 532 Indels. We kept all variants identified by *LRSomatic* with a VAF >0.05 with a quality score at or above 10 for PacBio data and 14 for ONT data. *ClairS-TO* indel calls on ONT tumor-only data however displayed a markedly different distribution in quality score than other ONT data (**Fig. S1**). To avoid excluding true indels, ONT tumor-only indel calls were hence filtered at a reduced quality threshold of 8.

In paired tumor-normal mode, *LRSomatic’s ClairS* module achieved F1 scores of 0.93 for SNVs and 0.69 for indels using PacBio HiFi data, and 0.91 for SNVs and 0.70 for indels using ONT data (**Fig. 5**). These results align with the reported performance for *ClairS* (0.95 F1 score for SNVs and 0.66 for indels in comparable datasets). As expected, tumor-only analyses showed slightly reduced performance, with F1 scores of 0.65 and 0.73 for SNVs and 0.23 and 0.27 for indels for PacBio and ONT, respectively (**Fig. 5**). These values are consistent with *ClairS-TO*’s reported sensitivity of 0.62 and 0.19 for SNVs and indels respectively. We anticipate the lower precision of tumor-only calling will benefit from future caller improvements and development of long read-based panels of normals for germline filtering. Taken together, these results demonstrate that *LRSomatic* supports state-of-the-art SNV and indel calling across long-read sequencing platforms in both tumor-normal and tumor-only mode.

**Figure 5:**
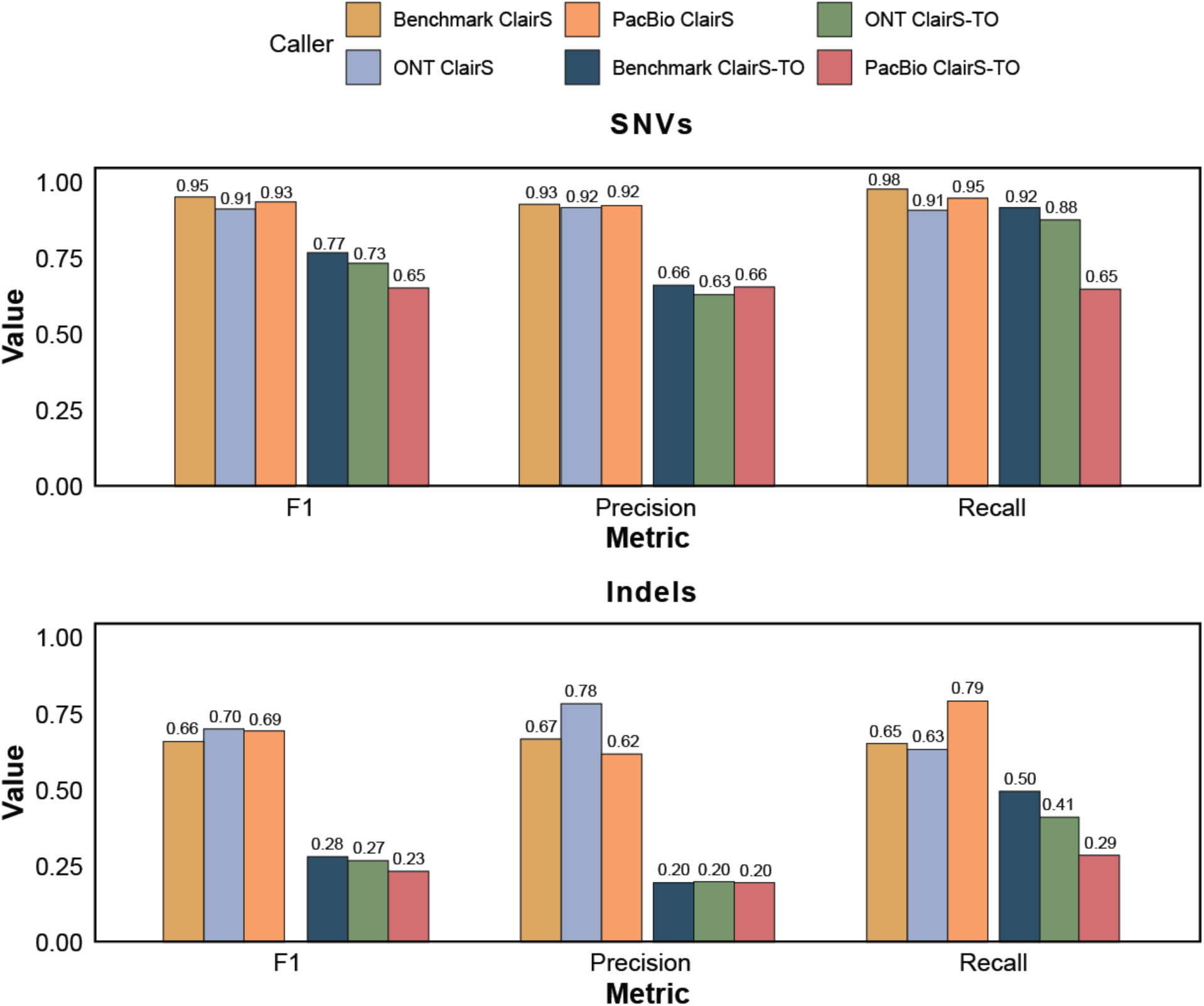
Summary of LRSomatic benchmarking on COLO829 paired and tumor-only runs with PacBio and ONT sequencing data. F1 scores, Precision, and Recall obtained for SNV (top) and Indels (bottom).

### Somatic Structural Variant Calling

To assess *LRSomatic’s* performance in detecting larger genomic rearrangements, we compared its *Severus* output for the HG008 and COLO829 samples in paired and tumor-only modes to established truth sets of somatic SV calls^43,47^. The COLO829 truth set contained 32 deletions, 13 translocations, 7 inversions, 7 duplications, and 3 insertions. The HG008 truth set contained 46 duplications, 42 deletions, 30 translocations, and 16 insertions. Keeping in line with prior *Severus* benchmarking, we defined SVs as sequence changes ≥50 base pairs and required that their VAF ≥0.1^33^. For comparison to the reference set, we allowed ≤500 bp discrepancy between the called positions of SV breakpoints^33^.

In paired tumor-normal mode, *LRSomatic* demonstrated robust SV detection capabilities. For COLO829, the workflow achieved F1 scores of 0.71 (PacBio) and 0.74 (ONT), while HG008 data yielded F1 scores of 0.91 (PacBio) and 0.74 (ONT) (**Fig. 6**). These results align with *Severus’* median reported performance of 0.88 in ONT data and 0.87 in PacBio, confirming that *LRSomatic* effectively identifies complex somatic structural variants including deletions, duplications, inversions, and translocations across both long-read platforms.

**Figure 6:**
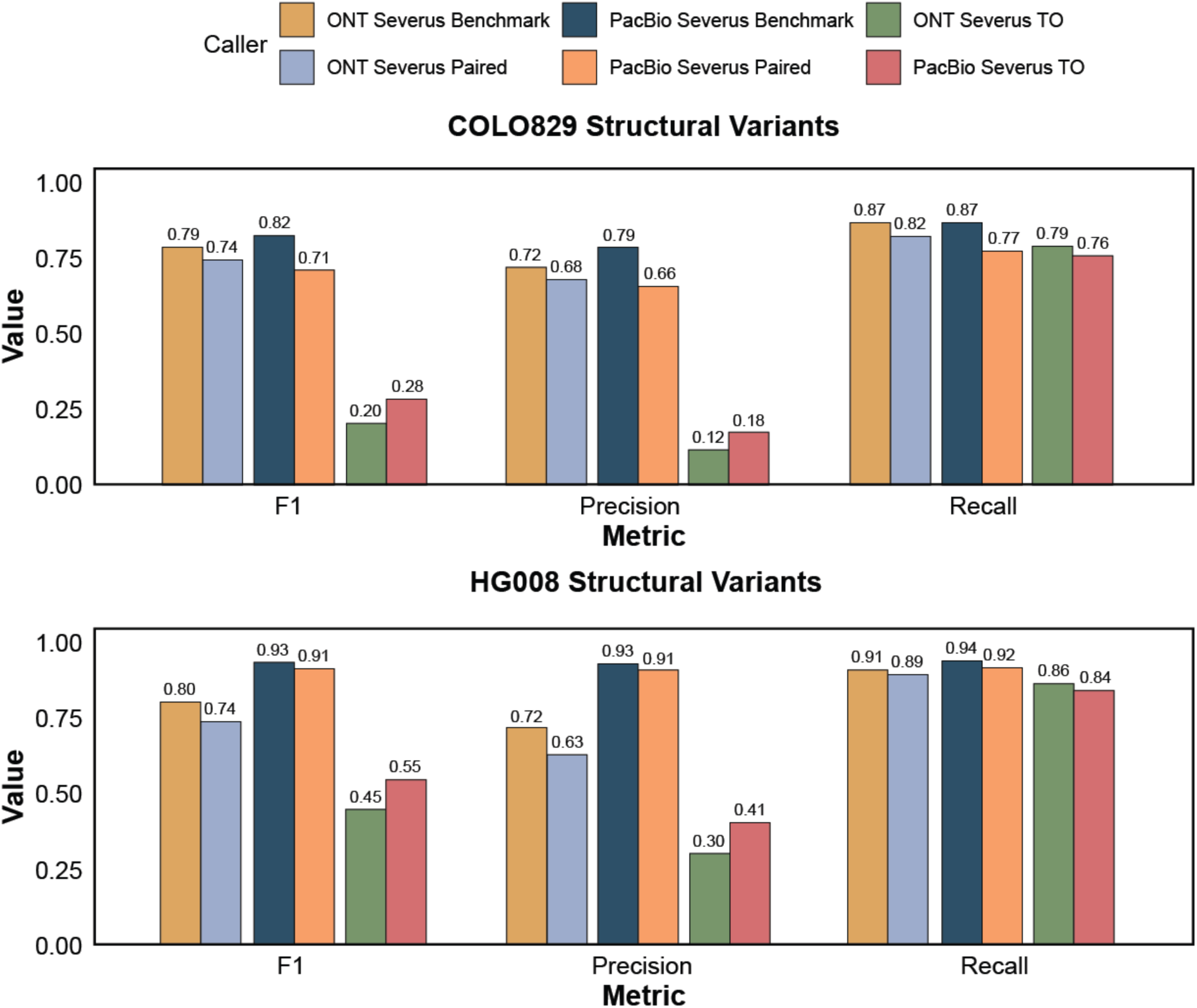
F1 score, precision, and recall for COLO829 (top) and HG008 (bottom) structural variants called in paired and tumor-only mode.

Tumor-only analyses showed reduced performance, with COLO829 achieving F1 scores of 0.20 (PacBio) and 0.28 (ONT), and HG008 yielding improved scores of 0.55 (PacBio) and 0.45 (ONT) (**Fig. 6**). As with the small variants, lower performance was primarily driven by decreased precision rather than sensitivity. This indicates that while *LRSomatic* successfully identifies true somatic SVs in tumor-only mode, it also reports a higher number of (likely rare) germline variants that cannot effectively be filtered without matched normal data given the current panel of normal references. Nevertheless, we show that *LRSomatic* provides state-of-the-art performance for detecting somatic structural variants.

### Clear Cell Sarcoma (CCS)

Finally, we assessed *LRSomatic’s* capabilities in a biologically and clinically relevant setting. Specifically, we performed PacBio and ONT-based Fiber-seq on metastatic and matched normal tissue from a case of sarcoma. CCS, specifically, is a rare form of sarcoma typically characterized by an *EWSR1*::*ATF1* translocation and resulting fusion transcript. We sought to determine if *LRSomatic* could identify clinically relevant somatic variant calls in this setting. The current gold standard workflow for comprehensive cancer analysis is *Oncoanalyser*, an *nf-core* pipeline for comprehensive reporting of variation in short read sequencing of cancer samples^45^. In its final output summary, *Oncoanalyser* reports key driver alterations along with copy number profiles and purity predictions^45^. In the absence of a comprehensive truth set for CCS15, we hence assessed whether *LRSomatic* recapitulates driver mutations identified by *Oncoanalyser* on corresponding short-read WGS data.

Specifically, *Oncoanalyser* identified the characteristic CCS fusion between *EWSR1* exon 8 and *ATF1* exon 4. In addition, it predicted three structural variants disrupting driver genes, including an inversion in *PMS2* and a deletion of *CDKN2A/B*. Moreover, the analysis found a 7 bp indel in *CHEK2* predicted to result in a frameshift. Finally, *Oncoanalyser* estimated a sample ploidy of 3.45 and a purity of 63%.

In both the PacBio paired and tumor-only data, *LRSomatic* called all reported somatic driver disruptions (**Fig. S2-5**). In addition, the analysis estimated the purity and ploidy at 81% and 3.43, respectively, with a copy number profile in line with *Onconalyser’s* (**Fig. 7** In both paired and tumor-only ONT sequencing data the *CDKN2A/B* deletion, inversion in *PMS2,* and most importantly the characteristic *EWSR1::ATF1* fusion were successfully identified. However, the *CHEK2* missense mutation was not identified in the paired and tumor-only ONT sequencing data, this may in part be due to lower coverage (19.90x vs 31.72x for ONT vs PacBio), as reads reporting the variants are visible upon inspection in IGV (**Fig. S2**). Finally, purity and ploidy were estimated at 5.09 and 65% respectively. The difference in ploidy here is due to an equivalent (doubled) solution being preferred by ASCAT, likely due to noise from the lower coverage ONT data (**Fig. S4, S6**). Taken together, this demonstrates *LRSomatic’s* ability to extract relevant somatic variation from long-read sequenced clinical samples.

**Figure 7:**
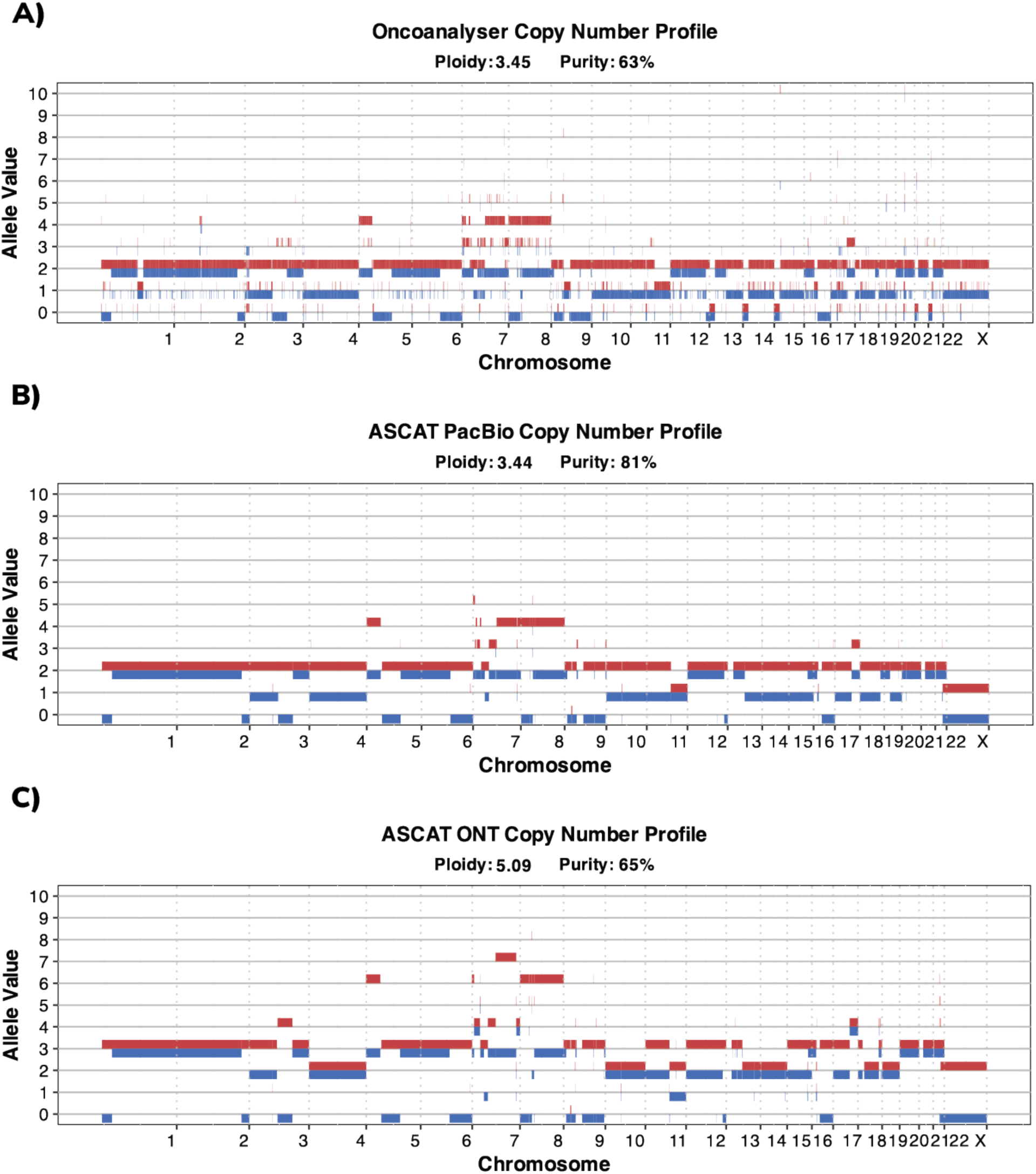
Allele-specific copy number profiles (major and minor alleles in red and blue, respectively), purity, and ploidy of clear cell sarcoma case CCS15 generated by (A) Oncoanalyser’s PURPLE toolkit on tumor-normal Illumina sequencing data and (B-C) LRSomatic’s ASCAT module on tumor-normal Fiber-seq data from (B) PacBio and (C) ONT.

Moreover, by extending traditional ONT and PacBio sequencing with Fiber-seq, *LRSomatic* can harness methylation calls to predict chromatin accessibility. For instance, the imprinted *MEG3* locus (*Maternally Expressed Gene 3*) shows clear haplotype-specific methylation (**Fig 8**). Haplotype one displays increased promoter methylation (5mC) and reduced accessibility (6mA labeling) compared to haplotype two. These observations likely reflect differential activity between the maternal and paternal allele and match expectations at imprinted genes. By harnessing Fiber-seq’s ability to capture nucleosome and chromatin accessibility patterns, *LRSomatic Fibertools* output boosts functional interpretation of somatic genomic alterations compared to standard WGS analysis.

**Figure 8:**
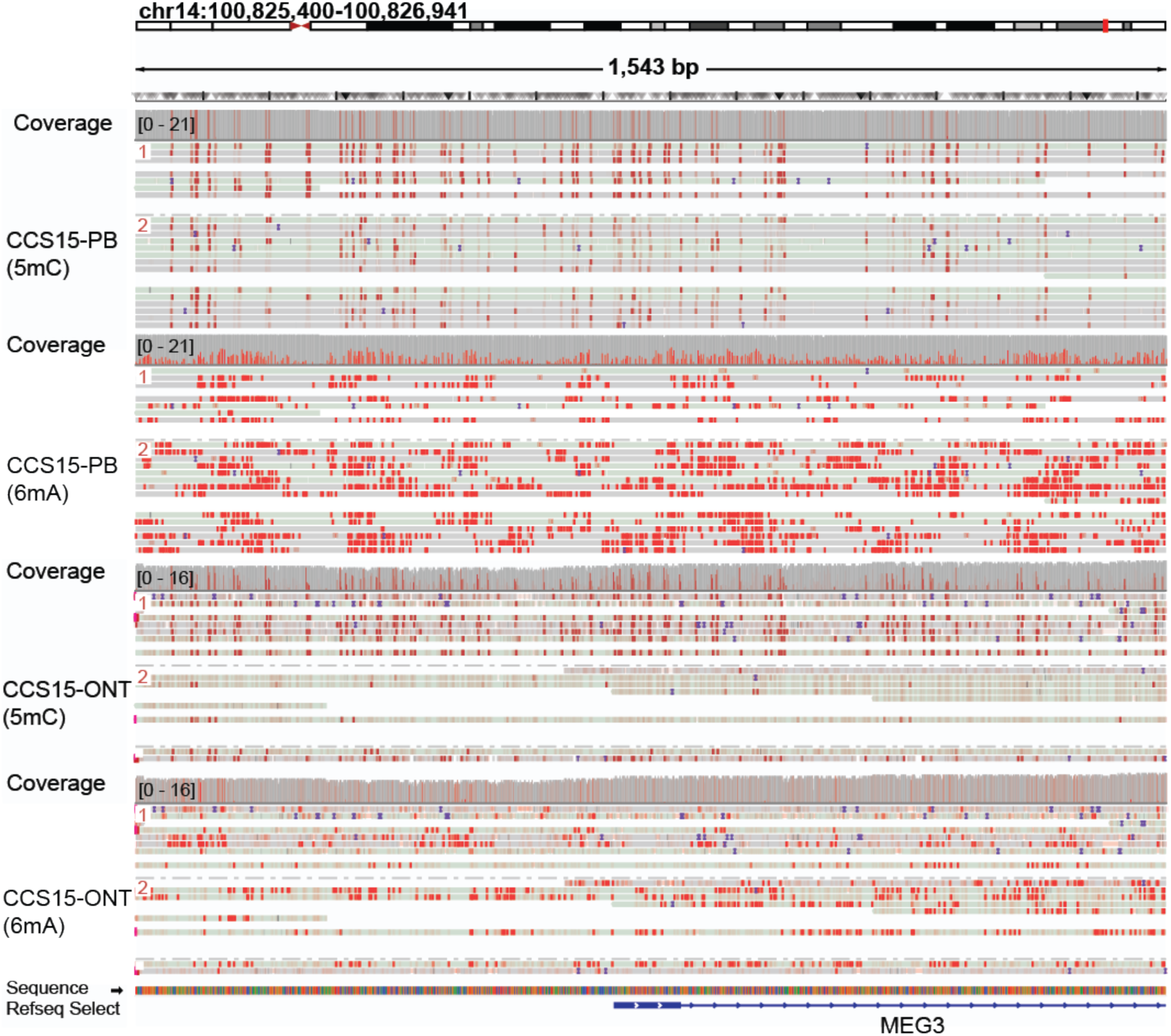
IGV screenshots of the imprinted MEG3 promotor region in PacBio and ONT Fiber-seq data. Both predicted positions of 5mC and 6mA modifications are displayed in red. Haplotype-specific differences can be seen in promoter methylation (5mC) and chromatin accessibility, reflecting distinct biological activities of the two alleles.

## Conclusions

Because of its superior ability to resolve complex genomic structure and variation, long-read sequencing was named Nature’s ‘Method of the Year’ in 2022^48^. Although the award was given in large part due to long-read sequencing’s key role in the construction of the first truly complete human reference genome, it has been critical in many discoveries since^14,48^. Long-read sequencing has been used to unveil novel SVs in cohorts of ovarian, colorectal, and breast cancers as well as to shine a light onto the mechanisms driving the creation of fusion genes in mucoepidermoid carcinoma^49–52^. Moreover, it has been used to resolve mechanisms by which the hepatitis B virus can drive relevant chromosomal rearrangements in hepatocellular carcinoma^53^. Outside of cancer, long reads have been used to reveal variation in highly repetitive regions in brains of patients with Alzheimer’s disease^54^. This resolving power will further our understanding of the role of complex variation in health and disease.

*LRSomatic* ties together state-of-the-art analysis tools to streamline comprehensive characterization of somatic variation from long-read sequenced samples. Established workflows for detecting somatic variation, such as *Oncoanalyser*, are designed exclusively for short-read data. However, short reads struggle to resolve complex variation, including SVs and variants in repetitive regions, a critical limitation when analyzing genomically complex samples or when more comprehensive analyses are desired. *LRSomatic* addresses this gap by enabling users to seamlessly process long-read data sets and obtain more complete profiles of somatic variation, regardless of sequencing platform or compute environment.

We demonstrate that *LRSomatic* accurately detects somatic variation across the full spectrum—from SNVs and indels to large SVs—in both PacBio and ONT sequencing data, using benchmarked COLO829 and HG008 datasets. Furthermore, we confirmed that *LRSomatic* recovers known epigenetic effects and clinically relevant somatic variants from a long-read sequenced clear cell sarcoma patient sample. While there remains significant germline bleed-through in tumor-only samples, we anticipate that this will be resolved by future long read-based additions to normal reference panels. Importantly, *LRSomatic* successfully identifies the vast majority of true somatic variants in the tested samples, demonstrating its robust performance as an analytical workflow for capturing somatic variation in long-read sequencing data.

## Supplementary information

Supplementary figures are available online.

## Code availability statement

The LRSomatic pipeline is available at https://github.com/intgenomicslab/lrsomatic. The code used to run all analyses is available at https://github.com/robert-a-forsyth/lr_somatic_paper_analysis.

## Author Contributions

Robert A. Forsyth: Methodology, Software, Visualization, Formal analysis, Data Curation, Writing – Original Draft. Luuk Harbers: Methodology, Software, Data Curation, Writing – Review & Editing. Amber Verhasselt: Software. Ana-Lucía Rocha Iraizós: Software. Sidi Yang: Software. Joris Vande Velde: Investigation. Christopher Davies: Investigation. Nischalan Pillay: Resources, Supervision. Laurens Lambrechts: Software, Writing – Review & Editing, Supervision. Jonas Demeulemeester: Conceptualization, Funding Acquisition, Resources, Project Administration, Supervision, Writing – Review & Editing.

## Acknowledgements

This work was supported by VIB and KU Leuven Internal Funds (C14/22/125 SymBioSys) and project grants from Stichting Tegen Kanker (2024-189 AGENDAS), the Research Foundation – Flanders (FWO; G021725N) and VIB (Grand Challenge GC06-C05). It was further enabled by large-scale research infrastructure funding from FWO (I014324N, T2T-Biology). The compute resources and services used in this work were provided by the VSC (Flemish Supercomputer Center), funded by FWO and the Flemish Government. We are grateful to the UCL/UCLH Biobank for Health and Disease (Royal National Orthopaedic Hospital satellite) for providing tissue for CCS15 and to the Edward Showler Foundation for generously supporting research related to Clear Cell Sarcoma. Robert Forsyth is supported by the Belgian American Educational Foundation. Ana-Lucía Rocha Iraizós is a doctoral fellow of the FWO (1129725N). Laurens Lambrechts is a postdoctoral fellow of Stichting Tegen Kanker (2025-031). We would additionally like to acknowledge Akanksha Farswan, Boyu Yu, the EMBL Protein Expression and Purification Core and the Leuven Genomics Core for their assistance and input.

