## Supplementary Figures for "LRSomatic: a highly scalable and robust pipeline for somatic variant calling in long-read sequencing data"

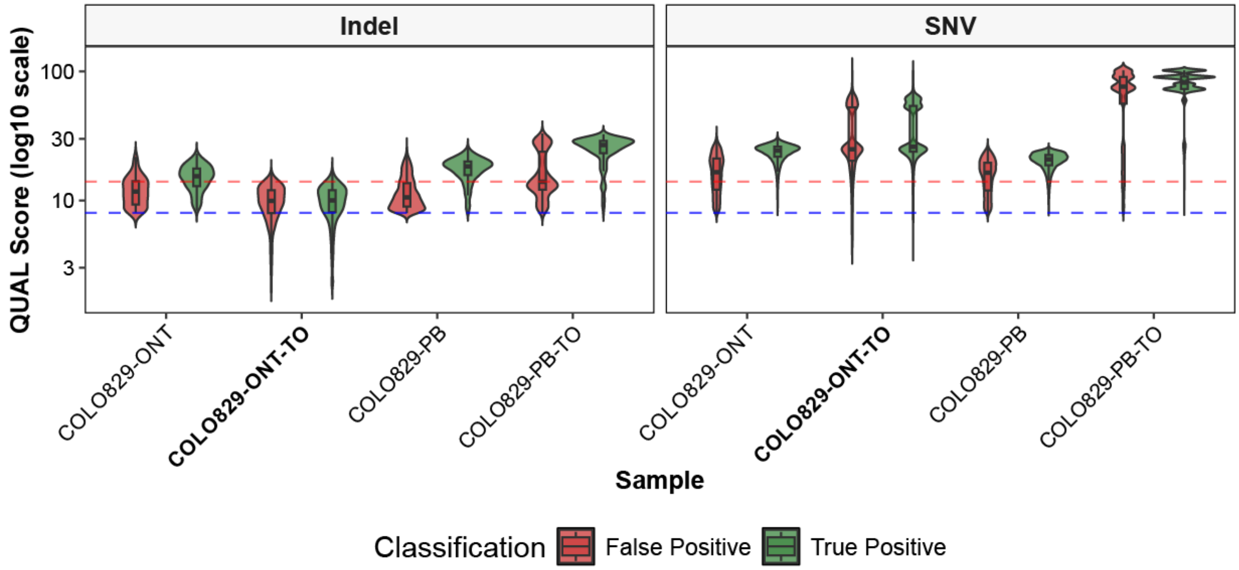


***Figure S1:*** *Distribution of quality scores in COLO829 SNV and indel call sets filtered at default ClairS and ClairS-TO quality thresholds (8), as opposed to our filtering thresholds (14 for all sets except ONT-TO indels). The distributions are colored by False Positive and True Positive according the COLO829 truth sets. Aside from COLO829-ONT tumor only indel calls, there are more false positive calls (with filtering threshold at quality score 8) with quality scores between 8 and 14 than true positive scores. For the COLO829-ONT-TO the majority of all indel calls have quality scores between 8 and 14, as a result we chose to keep the default filtering threshold for these data.*

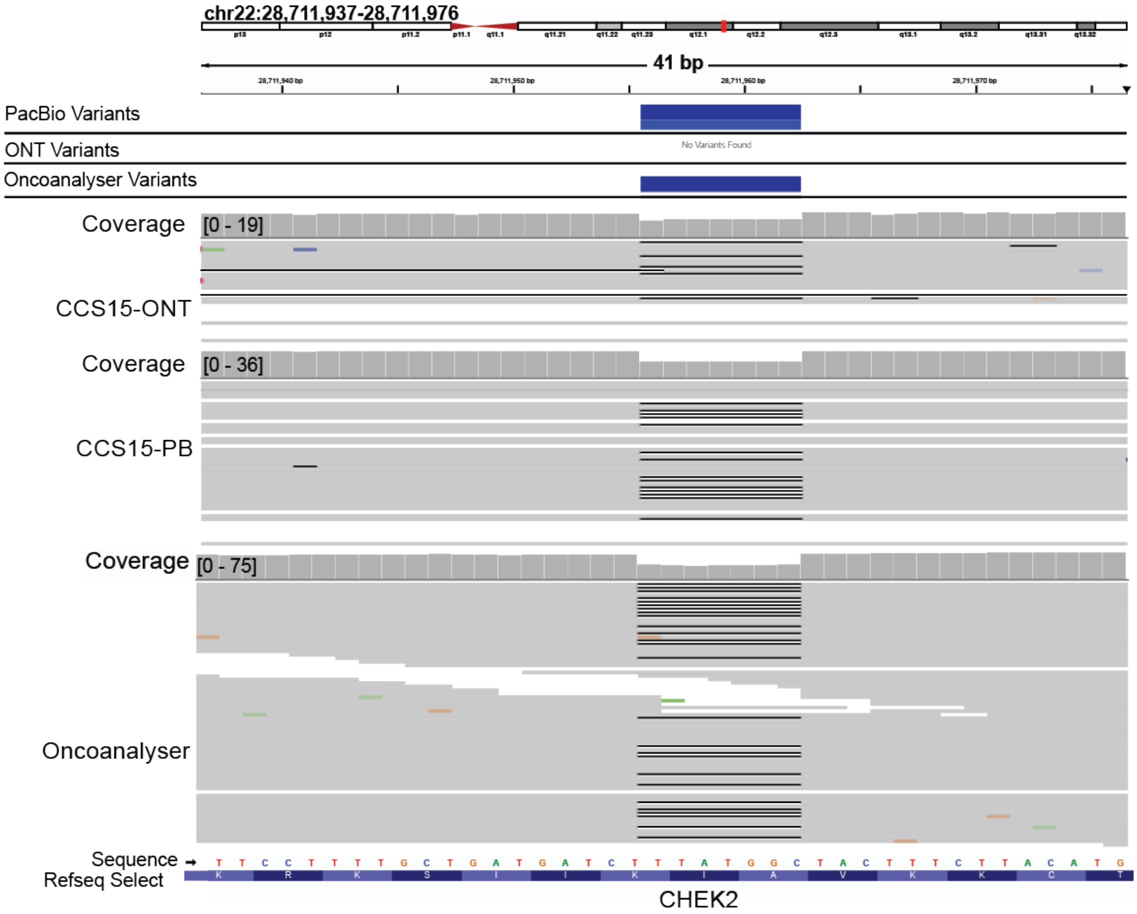


***Figure S2:*** *IGV screenshot highlighting the CCS15 CHEK2 p.A247fs driver variant. Tracks from top to bottom are: PacBio Somatic Variant Calls, ONT Somatic Variant Calls, Illumina small somatic variants called by Oncoanalyser, and aligned reads for PacBio, ONT and Illumina, respectively.*

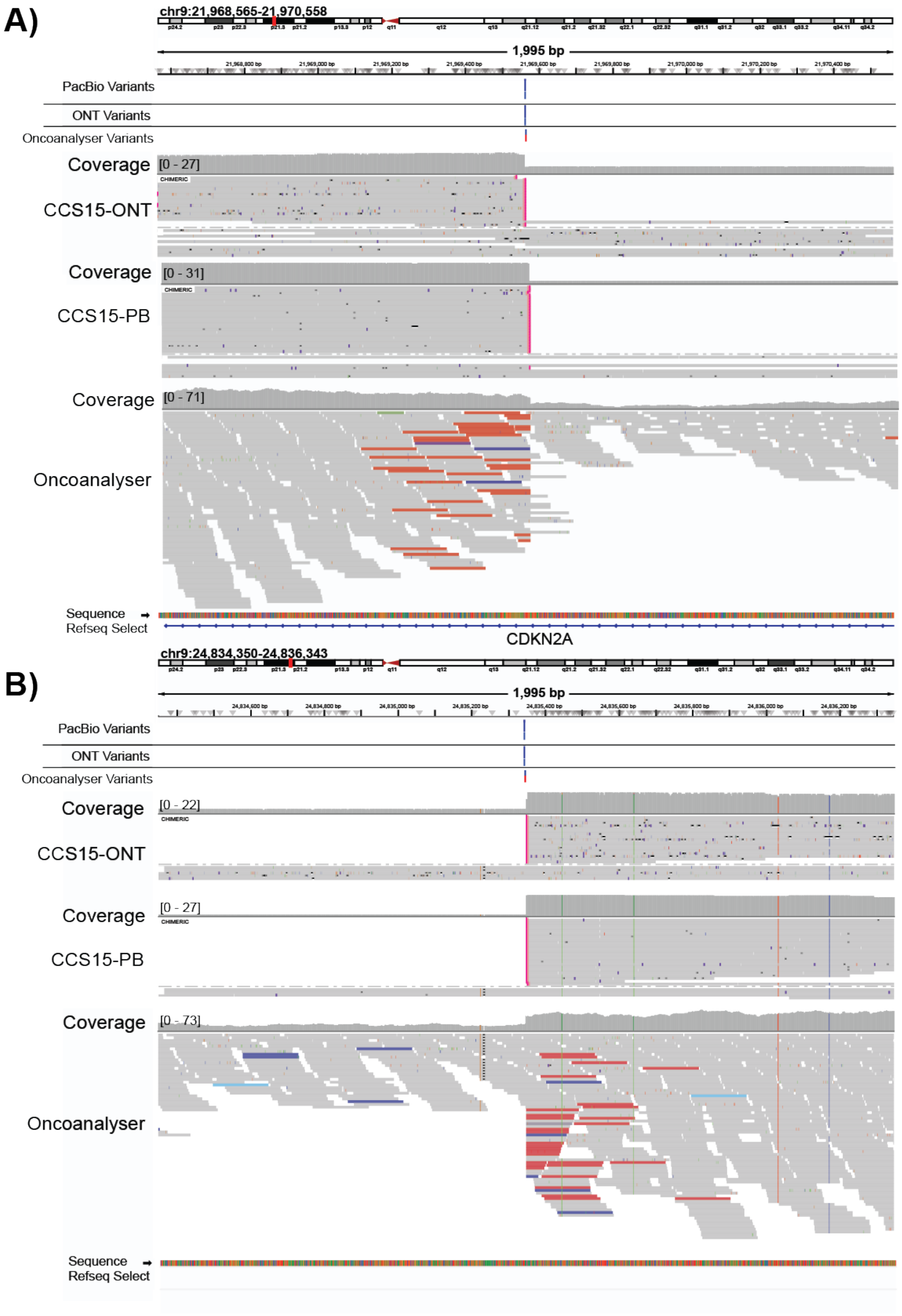


***Figure S3:*** *IGV screenshot highlighting the proximal (lower genomic coordinate) breakpoint* ***(A)*** *and the distal (higher genomic coordinate) breakpoint* ***(B)*** *CCS15 CDKN2A deletion. Tracks from top to bottom are: SVs for ONT, PacBio, and Illumina, and aligned reads for ONT, PacBio and Illumina, respectively for both* ***A*** *and* ***B****.*

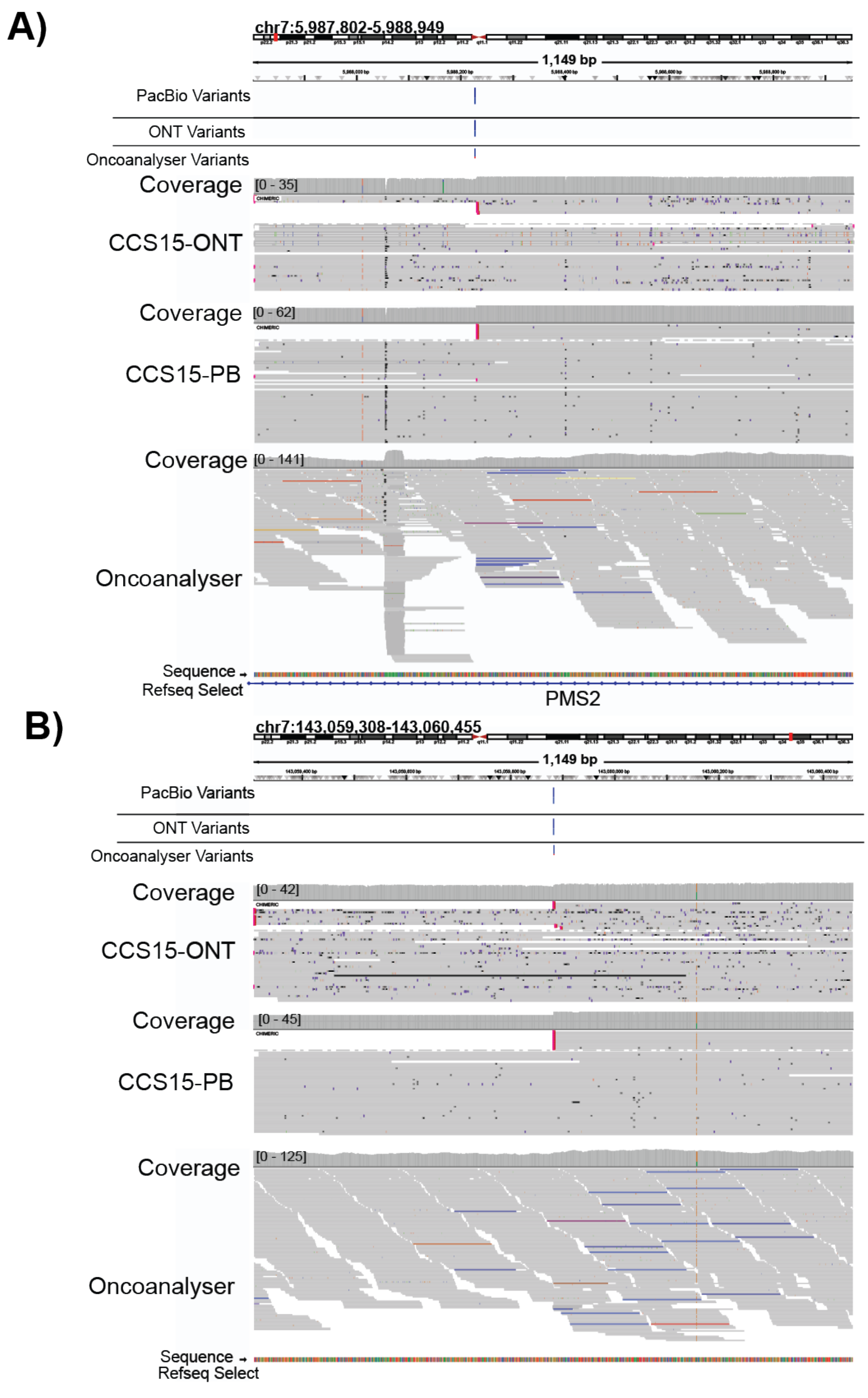


***Figure S4:*** *IGV screenshot highlighting the proximal (lower genomic coordinate) breakpoint* ***(A)*** *and the distal (higher genomic coordinate) breakpoint* ***(B)*** *the CCS15 PMS2 inversion. Tracks from top to bottom are: SVs for ONT, PacBio, and Illumina, and aligned reads for ONT, PacBio and Illumina, respectively for both* ***A*** *and* ***B****.*

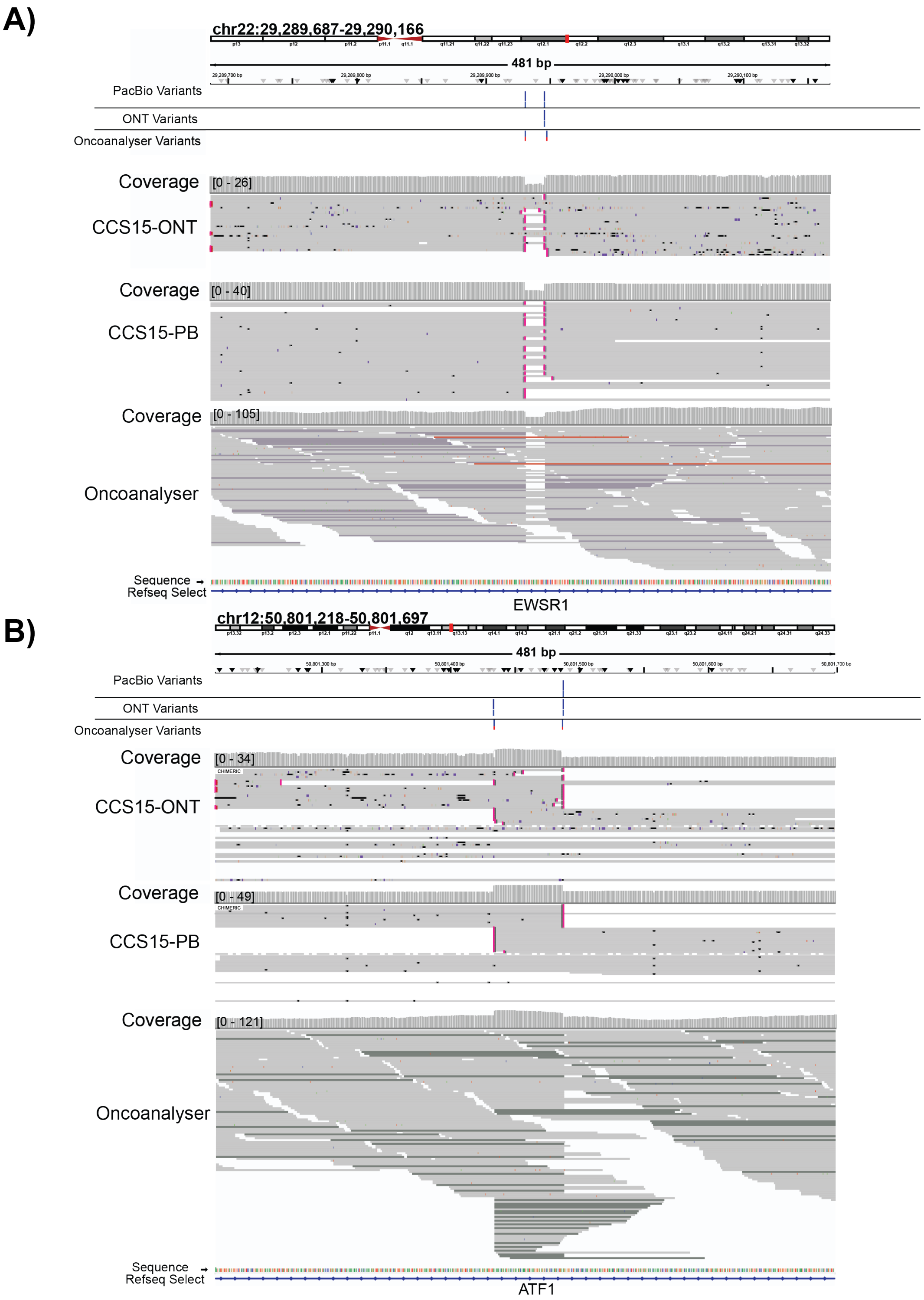


***Figure S5:*** *IGV screenshot highlighting the CCS15 characteristic EWSR1* ***(A)****-ATF1* ***(B)*** *fusion. Tracks from top to bottom are: SVs for ONT, PacBio, and Illumina SVs, and aligned reads for ONT, PacBio and Illumina, respectively for both* ***A*** *and* ***B****.*

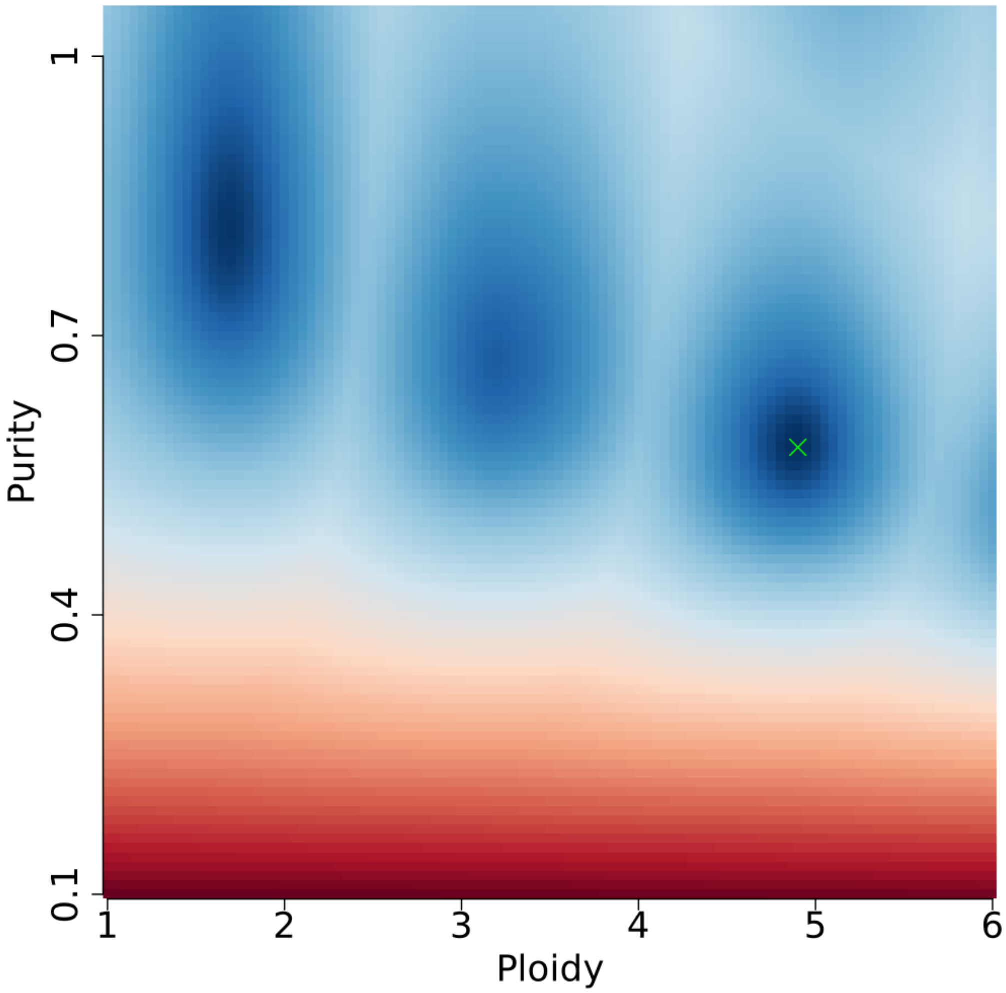


***Figure S6:*** *Sunrise plot for CCS15 ONT data displaying ASCAT's identified purity and ploidy fit scores, with red indicating poor solutions and blue indicating local optima. The final solution chosen is given by the green cross. A second optimum, corresponding to approximate ploidy and purity values of, respectively, 3.2 and 65%, is in agreement with the fits from Illumina and PacBio data.*
